# Decomposing the neural pathways mediating value-based choice

**DOI:** 10.1101/171744

**Authors:** Timothy R. Koscik, Vincent Man, Andrew Jahn, Christina H. Lee, William A. Cunningham

## Abstract

Understanding the neural implementation of value-based choice has been an important focus of neuroscience for several decades. Although a consensus has emerged regarding the brain regions involved, disagreement persists regarding precise regional functions and how value information flows between value-based choice regions.

In the current study, we isolate neural activity related to decision-making using a gambling task where expected gains and losses are dissociated from the received outcomes of choices. We apply multilevel modelling and mediation analysis to formally test whether brain regions identified as part of the value-based choice network mediate between perceptions of expected value and choices to take or pass a gamble.

A critical function in decision-making is accruing and representing value information to drive choice. Several regions have been assigned this role, including ventromedial prefrontal (vmPFC) and posterior parietal cortex (PPC), and the ventral striatum (VStr). The implied chain of events is one where regions that support the process of gathering relevant information mediate the relationship between choice and representations of value in other brain regions. Here, we formally test whether distinct brain regions express interregional mediation consistent with this chain of processes.

We observe that activity in vmPFC does not predict choice, but rather is highly associated with outcome evaluation. By contrast, both PPC and VStr (bilaterally) mediate between expected value and choice. Interregional mediation analyses reveal that VStr fully mediates between PPC and choice. Together these results suggest that VStr, and not vmPFC nor PPC, functions as an important driver of late stage choice.

**Significance Statement:** Making choices that maximize gain and minimize loss is critical for success. Our paradigm and analytic approach allowed isolation of choice-related neural signals from outcome-related signals. The vmPFC is involved at outcome rather than at choice. Isolating choice-related neural activity, we formally demonstrate that VStr and PPC mediate between expected value and choice. Our approach adds significant innovation by using generalized multilevel modelling to predict behavior with concurrent neural activity and formally testing the fully mediated pathway from stimulus through neural activity to behavior. Applying interregional multilevel mediation analysis, we demonstrate that ventral striatum comprises a final, critical step in processing value-based choice, mediating the relationship between value representation and choice.

## Introduction

Processing the relative value of options is critical for making appropriate choices. Research in decision neuroscience has consistently identified a network of brain regions associated with value computations, including: ventromedial prefrontal cortex (vmPFC), ventral striatum (VStr), posterior parietal cortex (PPC), anterior cingulate, amygdala, insula, as well as dorsomedial and dorsolateral prefrontal cortex (for recent reviews see 1, 2–10). Despite consensus on a general value-based choice network, questions remain regarding the individual contributions of each region. It remains unclear whether specific functions are ascribed to unique regions or distributed across multiple regions, and more importantly, how multifaceted choice-related information is integrated across the value-based network.

Because valued-based choice requires integration of multiple sources of information (e.g., costs, benefits, magnitudes, and probabilities), processes that weigh and aggregate value information are critical for choice. Activity in the PPC covaries with neural activity relating to value representation and motor output in other brain regions, and it is related to response times on decision tasks (11–13). This suggests that PPC aggregates information for choice and is more proximal to choice than processing elsewhere (Fig. 1A). However, PPC activity may be more related to numeric magnitude of options rather than value (14) consistent with localization of numeric representations (15). Other research highlights the vmPFC as a region that aggregates value information for choice, including representing chosen value at decision (16, 17), integrating value information (18), and being modulated by the duration of the decision-making process (19, 20). This suggests that vmPFC may take this interregional intermediary role (Fig. 1B). Likewise, the VStr is involved in dopamine neuromodulation of competing responses based on value (for a review see 21); given close relation to action, VStr may have an interregional intermediary role as well (Fig. 1C). These alternative predictions regarding the chain of processing events involved in value-based choice indicate that more research is needed to build toward a process model of value-based choice.

**Figure 1.**
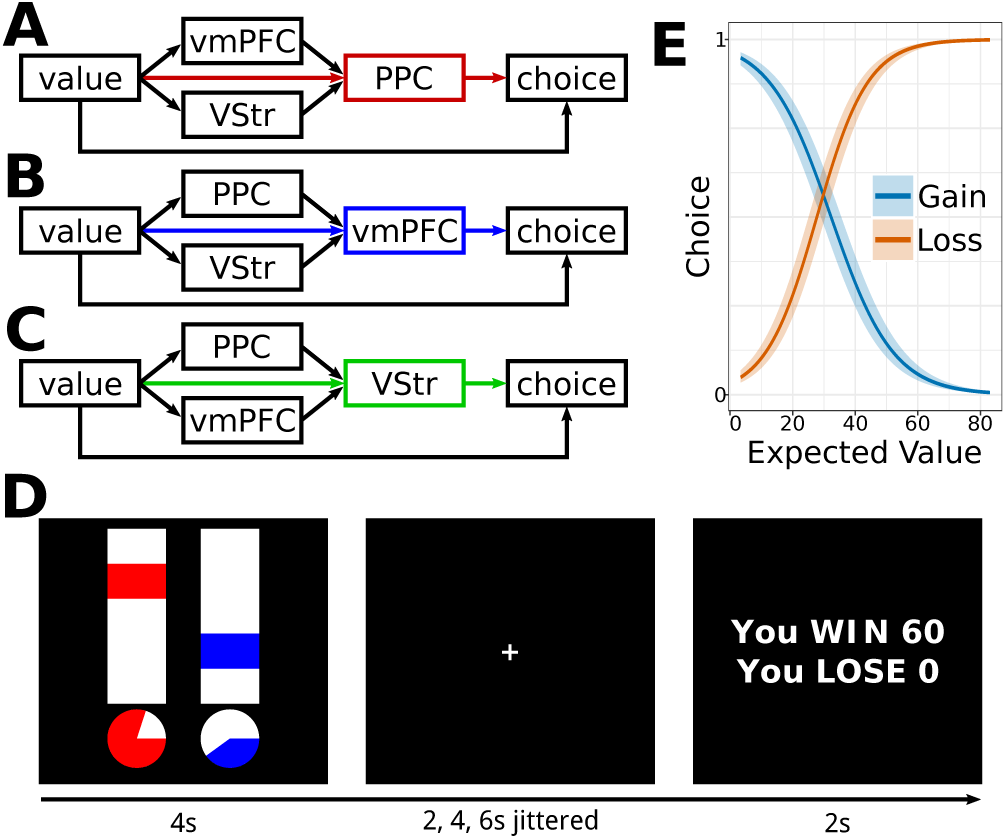
Alternative value-based choice network models: (A) PPC, (B) vmPFC, or (C) VStr are more proximal to choice than other regions. (D) Gamble task schematic (bars indicate points, pies indicate probability, color indicates gain or loss). Participants take or pass the gambles as a whole. (E) Participants tend to take gambles when the expected gain exceeds loss and pass when expected losses exceed gain.

The lack of clarity regarding these interregional relationships may stem from differences in experimental designs in decision neuroscience. For example, when outcome values are the same as expected values (e.g., one anticipates $10 and receives $10), it is difficult to determine whether neural activity reflects evaluation of expected value or evaluation of an inevitable outcome. These processes have distinct positions in the chain of events surrounding choice; decision processes precede choice and outcome evaluations follow choice. Confounding this temporal sequence not only limits the specificity of the functional roles that can be assigned to distinct brain regions, but also limits our ability to uncover how these regions operate together, and critically for the prediction of choice, which region assumes the most proximal value computation for choice. Differences between paradigms where expected and outcome values relate deterministically and those where they vary probabilistically offer clues to the potential function of value-based choice regions. For example, the vmPFC is more likely to be activated in studies where outcomes are deterministic (e.g., 11, 12, 18, 19, 22–27) relative to studies employing probabilistic outcomes (e.g., 28, 29), unless outcome-related signals, like prediction error or value at outcome, are being explored (e.g., 17, 30–32) (for exceptions see 13, 33–35).

Once expected and outcome values are dissociated, the chain of events in value-based decision is more clear: (a) stimuli that denote expected value of a choice are presented, (b) the brain processes this information and forms decisions, and (c) decisions are converted to motor programs that implement the choice. Brain regions mediate between perceptions of value and choice, and since expected values are experimentally manipulated, we can be certain about the direction of events. Once identified as value-choice mediators, the ordering of interregional relationships becomes critical. Among these regions, there is an implied chain of events specified by value representations, aggregation processes, and motor programs implementing choice. Interregional mediation analysis allows formal comparison of models that localize the most proximal aggregation of value for choice in distinct regions.

To better understand the value-based choice network, we measured brain activity while participants performed a gambling task. We adapted a duplex gamble task (36) where participants were presented with probabilistic gain and loss simultaneously; they chose whether or not to take each gamble (gain and loss as a whole) (Fig. 1D). To decompose activity specific to decision and outcome evaluation, gamble values were parametrically and temporally temporal distinct from outcome values on each experimental trial (see Methods for task details). This task design allows identification of regions involved at decision, at outcome, or involved in processing during both phases.

We test for relationships between neural activity, value, and choice using a multilevel modelling approach, where we can appropriately model the hierarchical structure of BOLD fMRI data (i.e., trials nested within scanning runs nested within participants), and thus model these distinct sources of variance (e.g., 37). We then extend the logic of the brain-as-predictor approach (38) into a voxelwise logistic multilevel framework to model choices as a direct product of concurrent neural activity and environmental parameters. A key difference between our study and previous studies is the voxelwise application of generalized multilevel models data instead of regions of interest (22).

Potential value-choice mediating regions must meet the following criteria: (a) activity must be predicted by expected value, and (b) activity must predict choice while controlling for expected value (for a thorough discussion of mediation see 39). We formally test the fully mediated pathway where neural activity mediates between expected values and choice. We are interested in trial-level mediation: that people tend to exhibit a particular response due to changes in neural activity on that trial instead person-level effects such as their membership in a condition or group. By applying multilevel mediation methods, we can formally explore whether neural activity mediates choices on a trial-by-trial basis (40, 41). Finally, once regions that mediate choice have been identified, interregional relationships can be tested, where trial-level activity in one region may mediate between activity in other regions and choice (Fig. 1A-C), providing insight into which regions provide the most proximal process in relation to choice.

## Results

Generalized multilevel modelling revealed that expected value of gains and losses significantly predict choices [gain: β=.156, z=21.62, p<.0001; loss: β=−.128, z=−21.24, p<.0001]. Participants tended to take gambles when expected gain exceeded loss and pass when expected loss exceeded gain (Fig. 1E).

### Neural Representations of Value at Decision and Outcome

To explore representations of value at the time of decision, we modelled single-trial beta estimates of neural activity for decision phases only. We applied voxelwise, linear multilevel modelling to predict brain activity with expected gain and loss values (see Methods and SI Methods). Upon request, we will provide access to all study materials and raw data hosted on our server for replication of our analyses and procedures.

Several regions are significantly predicted by both expected gain and loss value at decision, including: left PPC [(−32, −79, 30), size=24, expected gain: β=. 132, t=3.53, p<.001; expected loss: β=−. 147, t=−3.97, p<.0001], right VStr [(12, 8, −8), size=36, expected gain: β=. 133, t=4.03, p<.0001; expected loss: β=−.147, t=4.47, p<.0001] and the left Vstr [(−10, 8, −10), size=37, expected gain: β=. 148, t=3.7, p<.0001; expected loss: β=−.156, t=−3.93, p<.0001] (Fig. 2A). Additional regions that display this pattern of activity include a right lateral prefrontal region [rlPFC, (46, 38, 8), size=19, expected gain: β=. 137, t=3.59, p<.001; expected loss: β=−.119, t=−3.14, p<.01] and a region of left pre- and postcentral gyrus [pCG, (−30, −28, 60), size=111, expected gain: β=.147, t=4.08, p<.0001; expected loss: β=−.137, t=−3.83, p<.001]. In these regions, activity increased with increases in expected gain and decreased with increased expected loss (Fig. 2B). By contrast, a region of visual cortex [(0, −88, 12), 313 voxels] increased activity to both expected gain [β=.273, t=5.49, p<.0001] and loss [β=.171, t=3.47, p<.001], consistent with increased looking or attention to extreme expected values (see Fig. S1 and Table S1 for full results).

**Figure 2.**
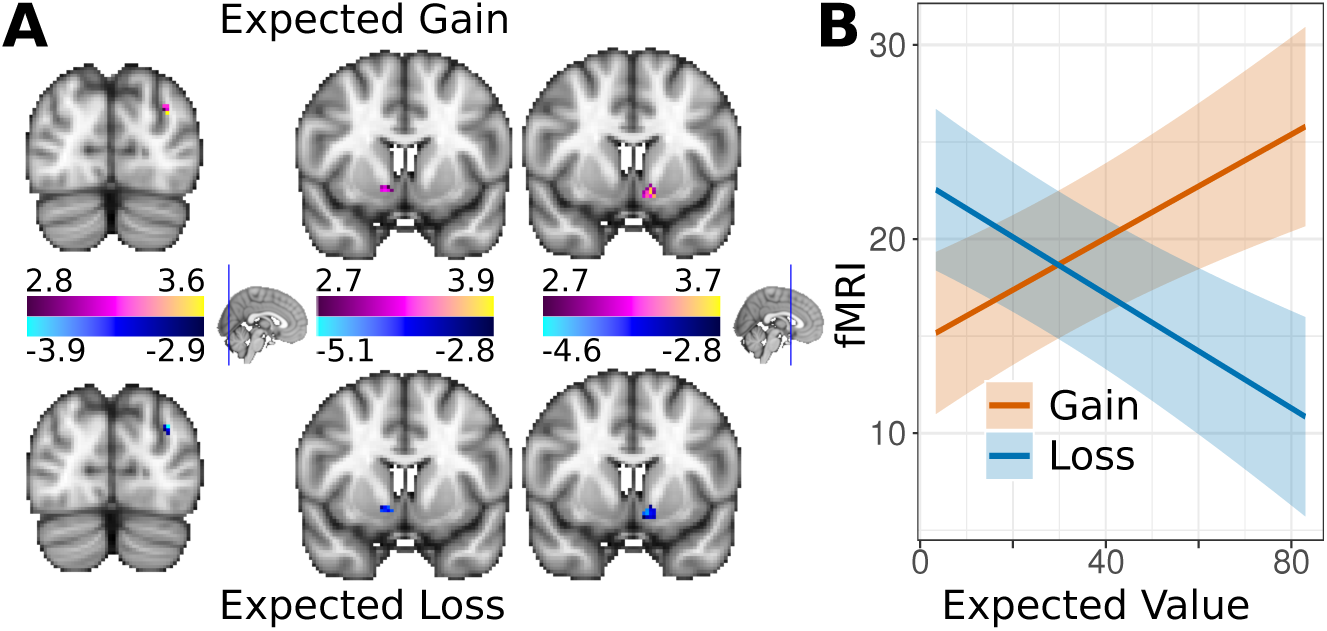
Neural Representations of Expected Values. (A) PPC and bilateral VStr (left to right) are predicted by both expected gain and loss. PPC and VStr increase activity when expected gain is greater and decrease when expected loss is greater (B, right VStr shown). Shaded regions indicate 95% confidence limits.

To explore whether decision-making regions with outcome evaluation regions, we modelled neural activity at outcome with both gain and loss values and choices (see Methods and Fig. S2 and Table S2 for full main effects and Fig. S3 and Table S3 for interaction effects). Activity in bilateral PPC (Fig. 3A) increased with both gain [right PPC: β=.244, t=7.35, p<.0001; left PPC: β=.175, t=6.1, p<.0001] and loss [right PPC: β=.125, t=3.98, p<.0001; left PPC: β=.12, t=4.38, p<.0001] at outcome (Fig. 3B). Activity in PPC did not distinguish outcome value, encoding magnitude not value. Interactions between gain or loss and choice were not significant in PPC.

**Figure 3.**
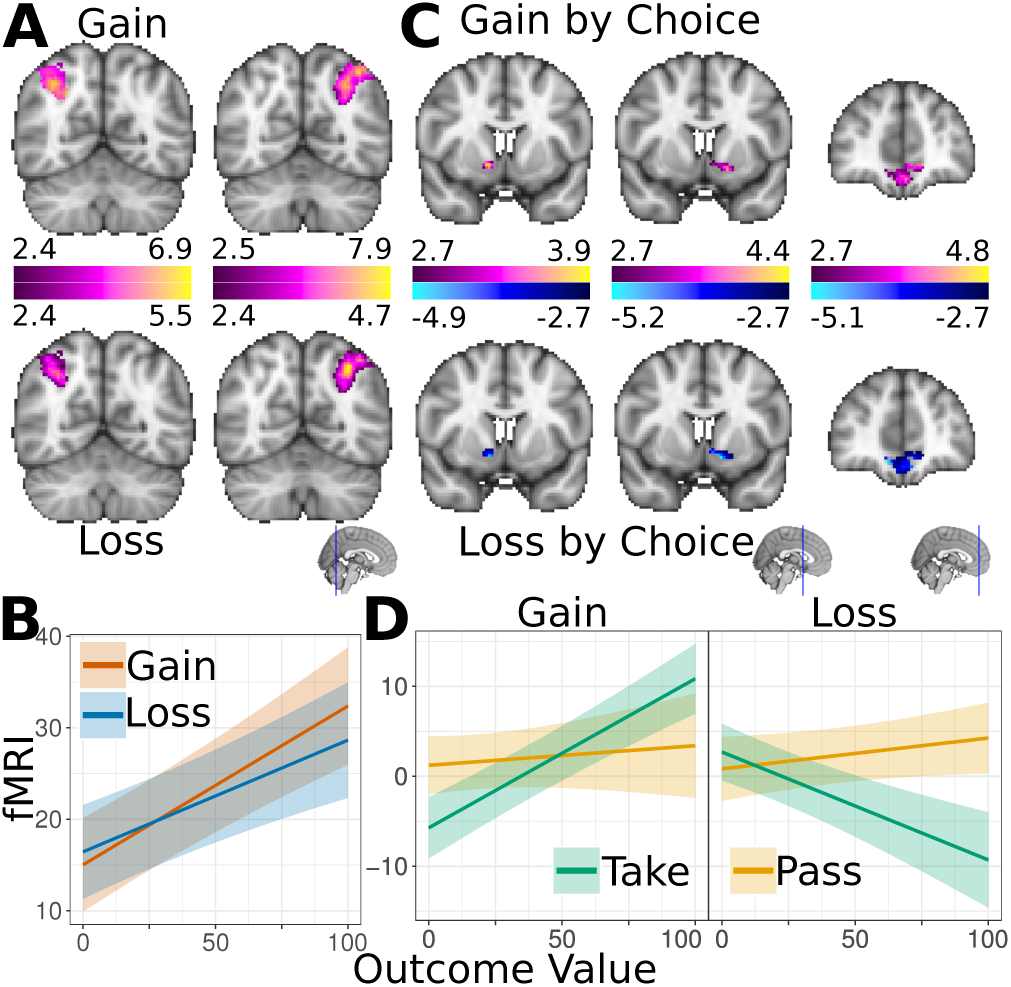
Neural Representations of Outcome Value. Linear multilevel models of outcome-related estimates of neural activity reveal that (A) PPC activity tracks gain and loss magnitude, irrespective of whether they were received or not (B). Activity in bilateral VStr and vmPFC (C) is related to the interactions between gain or loss and choice; increased with received but not missed gains and decreased with received but not avoided losses (D). Shaded regions indicate 95% confidence limits.

Activity in bilateral VStr and a region of vmPFC (Fig. 3C) were significantly predicted by the interaction between gain and choice [right VStr: β=.072, t=3.87, p<.001; left VStr: β=.082, t=4.39, p<.0001; vmPFC: β=. 118, t=4.32, p<.0001] as well as loss and choice [right VStr: β=−.077, t=−4.37, p<.0001; left VStr: β=−.083, t=−4.65, p<.0001; vmPFC: β=−.119, t=−4.58, p<.0001] at outcome. These regions encode outcome value, such that received gain and loss result in increased or decreased activity respectively, but avoided gain and loss are not strongly encoded (Fig. 3D).

VmPFC represents received gain and loss at outcome but not expected gain and loss at decision. Exploring vmPFC at decision, expected gain is not represented [β=.069, t = 1.53, p=0.13] but expected loss is [β=−.098, t=−2.18, p=.03]. Furthermore, trial-related activity in vmPFC does not predict choice at decision [β=−.001, t=−0.63, p=0.53], suggesting that it is not the most proximal value aggregation for choice (Fig. 1B), indeed vmPFC does not mediate between value and choice (IE=−.002, 95% CI =−.009−.003)].

### Identification of Neural Regions for Mediation Analysis

Potential value-choice mediating regions must (a) be predicted by expected value at decision, as demonstrated above, and (b) activity in these regions must also predict choice when controlling for expected value. We calculated voxelwise, logistic multilevel models where choice was predicted with trial-related neural activity while controlling for subject-related and scanning run-related activity and expected gain and loss (Fig. S4 and Table S4). Several clusters meeting these criteria were observed: bilateral VStr [right: (12, 6, −8), size=48; left: (−10, 8, −10), size = 48], and left PPC [(−32, −74, 28), size=12) as predicted (Fig. 4A), as well as pCG [(−30, 28, 60), size=132], rlPFC [(46, 38, 8), size=20], and posterior calcarine sulcus [(−2, −96, 6), size=27].

**Figure 4.**
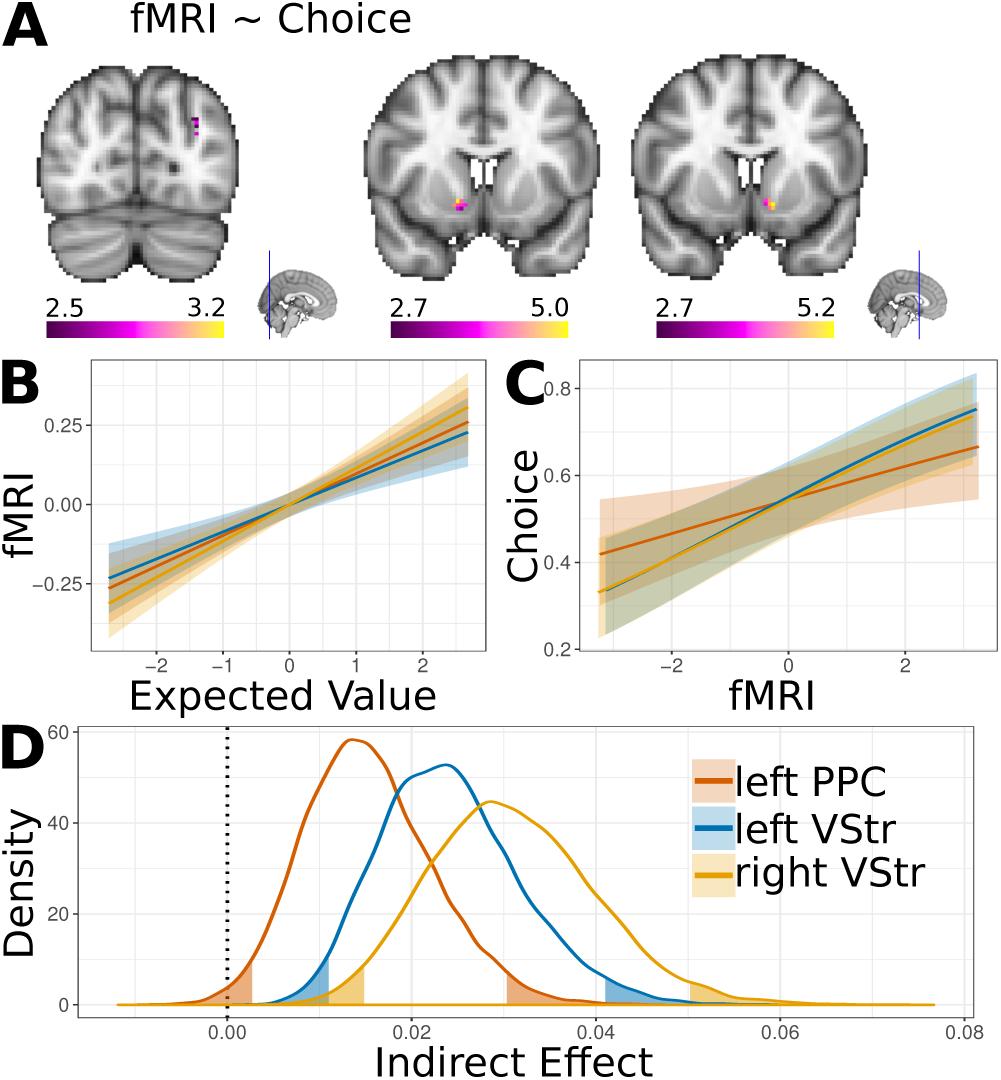
Neural Activity Predicting and Mediating Choice. Generalized multilevel models reveal voxels where trial-related activity predicts choice. (A) Regions include left PPC and bilateral VStr. (B) Activity in these regions is predicted by expected value, and (C) activity predicts choice. (D) Mediation analysis reveals that the indirect effect for each of these regions is significant and they mediate between expected value and choice. Shaded regions indicate 95% confidence limits or the inverse for density plots.

### Ventral Striatum and PPC Mediate between Expected Value and Choice

Within potential, value-choice mediating regions, we calculated the mean time series for each region, then applied a linear multilevel model, where expected value (expected gain - loss) predicted activity, and a generalized multilevel model, where choices were predicted by trial-related activity. We then utilized *stacked multilevel modelling* to estimate the covariance structure of these models simultaneously. Given the nested structure in multilevel data, stacked multilevel modelling accounts for the covariance of slopes in the indirect path (e.g., value to brain and brain to choice) in addition to modelling the variance in slopes for both components of the indirect path (41). This covariance structure was used to calculate the indirect effect (IE) and confidence intervals with Monte Carlo multivariate simulation (42).

Within bilateral Vstr, activity related to expected value [right VStr: β=.115, t=5.96, p<.0001; left VStr: β=.085, t=4.41, p<.0001] (Fig. 4B). Consistent with voxelwise models, logistic multilevel modelling of choices, trial-related changes in the mean time series in bilateral VStr predicted choices [right VStr: β=.27, z=4.25, p<.0001; left VStr: β=.283, p=4.67, p<.0001] (Fig. 4C). Thus bilateral VStr meets criteria for potential mediation. Mediation analysis reveals that activity in bilateral VStr significantly mediates the relationship between expected value and choice [right VStr: indirect effect (IE) =.031, 95% CI=.015−.05; left VStr; IE=.024, 95% CI=.011−.041] (Fig. 4D).

Within the PPC (Fig. 4A), activity was related to expected value [β=.097, t=5.03, p<.0001] (Fig. 4B) and activity predicted choice [β=.157, z=2.45, p<.01] (Fig. 4C). Similar to bilateral VStr, PPC significantly mediates the relationship between expected value and choice [right VStr: IE=.022, 95% CI=.008−.039] (Fig. 4D).

Two other clusters, pCG and rlPFC both relate to expected value [pCG: β=. 102, t=5.31, p<.0001; rlPFC: β=.087, t=4.47, p<.0001], activity predicts choice [pCG: β=.3, z=4.69, p<.0001; rlPFC: β=.242, z=3.86, p<.001], and activity mediates the relationship between expected value and choice [pcG: IE=.031, 95% CI=.008−.037; rlPFC: IE=.021, 95% CI=.008−.037].

Given the highly correlated nature of neural activity between regions, it is important to demonstrate that mediation is not the spurious result of common variance. To test this possibility, we explored the region within the visual cortex where activity increased in relation to expected gain and loss [but not combined expected value: β=.02, t=1.04, p=0.299], activity predicted choice [β=.233, z=2.78, p<.001], and activity was highly correlated with activity in other regions (left VStr: r = 0.39; right VStr: r = 0.41; and PPC: r = 0.41; all p’s < 0.001). The indirect path through visual cortex was not significant [IE=.031, 95% CI=−.015−.05] (Fig. S5). Thus, it is unlikely that significant mediation is due to some general, shared variance component of neural activity.

Motor processing in the brain is necessarily proximal to the actions that implement choice relative to value processes, e.g., motor cortex directly evokes muscle movements to enact choice. Thus, there motor cortex and action selection regions, such as the dorsal striatum (21) should mediate between value processes and choice. Indeed, we observed a region in left pCG (contralateral to right-handed responses) that mediates between value and choice. While task design features allow disambiguation of processes up to abstract representations of value-based choice, action-value representations were not conditional on any feature of the gamble task. Thus, these processes could not be distinguished here.

### Interregional Mediation in the Value-based Choice Network

Since multiple brain regions mediate between expected value and choice, it is important to understand how information flows between them in service of choice. We tested whether these value-choice mediators also mediate between the neural activity in other mediator regions and choice. VmpFC does not predict decision and has already been excluded as a mediator; the remaining alternatives regarding interregional mediation of choice, include the PPC (Fig. 1A) and VStr (Fig. 1C). A third alternative is that they function together to aggregate value information, where there is partial mediation for both pathways. We performed pairwise comparisons between value-choice mediators in both directions, e.g., whether right VStr mediated between left PPC and choice and whether left PPC mediated between right VStr and choice.

### Ventral Striatum Mediates the Relationship between PPC and Choice

First, linear multilevel modelling, where brain activity in one region is used to predict activity in a potentially mediating region, reveals that all pairwise comparisons between regions are significant, which is to be expected with highly correlated brain signals [all p’s<.0001] (Fig. 5A).

**Figure 5.**
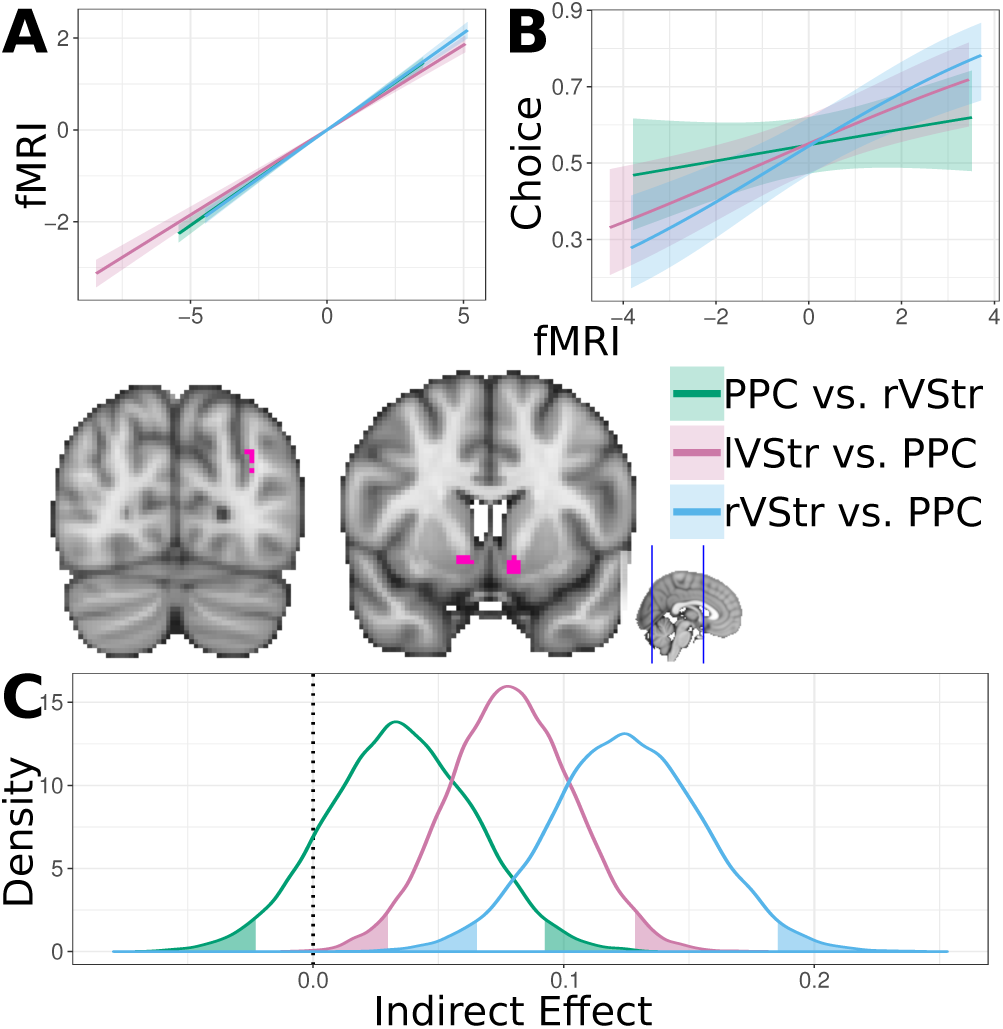
Interregional Mediation, Proximal Processing Regions for Choice. Activity in left PPC and bilateral VStr predicts activity in each of the other regions (A, *green:* right VStr predicting left PPC, *purple:* left PPC predicting left VStr, and *blue:* left PPC predicting right VStr). (B) Left PPC does not predict choice after controlling for either VStr (*green*, controlling for right VStr). Bilateral VStr predicts choice after controlling for PPC (*purple:* left VStr, *blue:* right VStr). (C) Moreover, bilateral VStr fully mediates between left PPC and choice (*purple:* left VStr, *blue:* right VStr), and left PPC does not mediate between either VStr and choice (*green*, PPC and right VStr shown). Shaded regions indicate 95% confidence limits or the inverse for density plots.

Second, generalized multilevel modelling predicted choices with brain activity in the potential mediating region while controlling for activity in the other brain region and expected value (Fig. 5B). Activity in right VStr predicted choice consistently [controlling for left VStr: (β=.212, x=2.2, p<.05; controlling for left PPC: β=.296, z=4.18, p<.0001], as did left VStr [controlling for PPC: β=.212, p<.01] except when controlling for right VStr [β=. 122, z=.13, p=.186]. Activity in PPC did not predict choice consistently [controlling for right VStr: β=.084, z=.12, p=.232; controlling for left VStr: β=.084, z=1.21, p=.228]. Among other regions, left pCG and rlPFC both predict choice after controlling for other regions (all p’s<0.036).

Finally, pairwise mediation analysis (Fig. 5C) indicated that bilateral VStr fully mediates between activity in the PPC and choice [right VStr: IE=.125, 95% CI=.067−.185; left VStr: β=.079, 95% CI=.029−.128]. By contrast, this flow of information is unidirectional for choice, where PPC does not mediate between right VStr [IE=.035, 95% CI=−.023−.093] or left VStr [IE=.03, 95% CI=−.019−.078] and choice. The results are consistent with the notion that bilateral VStr forms a pathway through which information must flow before making a decision (Fig. 1C). We also observe that left pCG [IE=.101, 95% CI=.04−.163] and rlPFC [IE=. 101, 95% CI=.054−.149] mediate between PPC and choice. Remaining pairwise mediation relationships, including those in motor cortex, appear to be partial (i.e., information flows bidirectionally), and mediation pathways through right lateral PFC were only present at the lowest (95%) thresholds (all other mediation effects reported are significant at 99% and 99% thresholds as well).

## Discussion

Understanding value-based choice requires understanding how stimulus values are manipulated by the brain to produce choice. Research in this regard has heretofore predicted neural activity from choices and/or task parameters. Using generalized multilevel modelling and mediation analyses, we provide the first direct test of how activity in reward-related brain regions mediates between expected value and choice. This mediation constitutes direct evidence that bilateral VStr and PPC transforms incoming information regarding the expected value of a decision to produce choices of whether to take a gamble.

Despite the literature regarding the role of the vmPFC as a value aggregator for choice (e.g., 17), our results indicate that vmPFC is unlikely to be the most proximal representations for choice. Our results are more consistent with a role for vmPFC in outcome evaluation rather than at decision. Moreover, activity in vmPFC voxels did not predict choice, thus it cannot take this proximal role at least for take/pass choices or the types of gain and loss in our task. Our data cannot preclude the possibility that vmPFC takes this role under other circumstances, e.g., comparing options or values vary in abstractness (food vs. money vs. points).

Our data suggests that the VStr and not the PPC is the most proximal value computation for choice. After accounting for VStr activity, the relationship between PPC and choice is eliminated, suggesting that the VStr fully mediates the pathway. Moreover, PPC is associated with gain and loss magnitude not value at outcome, which is consistent with prior research targeting this distinction (14) and reports regarding numeracy in PPC (e.g., 15). However, this specific interpretation is limited by the current study design, as numeric magnitude and monetary value were confounded. These results conflict with the observation that PPC activity relates to between-subjects differences in reaction time and activity in other value processing regions (11–13). In regard to decision-making, between-subjects differences in response time is not the correct level of analysis. Rather, trial-level changes in response times within-subjects reflect the aggregation processes necessary for decision-making. Moreover, previous analyses of relationships between PPC and value ignored their ‘causal’ direction. Future research could benefit from targeting trial-level differences in response time and apply the multilevel mediation methods demonstrated here to probe the direction of interregional relationships.

Evidence of significant mediation is insufficient to determine a causal relationship, but can act to rule out models that are causally less plausible. We began comparing three potential causal models where PPC, VStr, and vmPFC were considered as the most proximal value representation for choice. While the plausibility of the vmPFC in this role is diminished as it appears circumscribed for outcome rather than decision, mediation models indicate that the VStr is the most plausible of these alternatives. Of course, more data is necessary to make truly causal claims. This VStr function is consistent with the view that dopamine activity in the VStr influences action selection and modulation of choice behaviours in conjunction with the dorsal striatum (21, 43). Furthermore, our data are consistent with the notion that the VStr is a necessary common path between cortical and limbic value processing and the motor system (44, 45) including influences on cortical motor systems and potential direct (non-cortical) influences on locomotor activity via midbrain regions (46–49). Clarifying what this role is or what the relative contributions of dorsal and ventral striatum are will require further data. It is possible that the VStr is involved in evaluating all stimuli regardless of choice, or it might be inextricably linked to implementing value-based choice and active only when value-based choices are required. Future, hypothesis-driven research could leverage these multilevel mediation techniques to further elucidate the role of the VStr in relation to value-based decision-making and illuminate its role in relation to action selection and motor output regions.

## Conclusion

Our results provide a formal test that demonstrates that neural activity in the VStr and PPC mediate the relationship between expected value and choice. Moreover, the VStr provides a final common path between neural representations of value and choice. In addition, we provide an application of linear and generalized multilevel modelling to functional neuroimaging to account for the hierarchically structured error inherent to functional neuroimaging, and demonstrate that mediation models can provide evidence consistent with causal interpretations of brain activation.

## Methods

### Participants

We recruited 23 individuals (14 women; one woman was excluded for being unable to complete the task, 22 individuals were used) from the Queen’s University community through advertisements. Participants provided consent in accordance with the Queen’s University Institutional Review Board. All participants had no history of neurological or psychiatric disorder, had normal or corrected vision, and were right-handed. Participants were compensated with $40.

### Procedure

Participants performed a 10-minute pre-scanning task to familiarize them with the gambles to be used in the fMRI scanner. Gambles were presented as color-coded bar charts (red or blue) representing gain and loss magnitude (20, 40, 60, 80 points as bar height) and pie charts representing probabilities (13, 33, 50, 67, 83%) (Fig. 2). Color and position of gambles (gains on left or right) were counterbalanced across participants. Following a 2s fixation cross, gambles were displayed for 4 seconds during which participants made a button press to take or pass. Following each decision and a jittered delay of 2, 4, or 6s, feedback appeared for 2s indicating how many points had been (or could have been) gained and lost; both, either, or neither the gain and/or/nor loss could occur.

### fMRI Acquisition

Images were acquired with a Siemens Tim Trio 3T scanner. For whole-brain functional coverage, 32 axial slices (slice thickness=3.5mm, 0.5mm skip) were prescribed parallel to the AC-PC line. Nearly isotropic functional images were acquired from inferior to superior using a single-shot gradient echo planar pulse sequence (TE=25ms, TR=2s, in-plane resolution=3.5×3.5mm, matrix size: 64×64, and FOV=224mm).

### fMRI Preprocessing

fMRI scans were preprocessed using FSL (50) and included the following operations: (1) motion correction; (2) non-brain voxel removal; (3) spatial smoothing, 5mm FWHM Gaussian kernel; (4) intensity normalization using 4D grand mean; (5) temporal filtering, high pass Gaussian-weighted least squares straight line fitting (sigma=70s); and (6) affine registration to MNI standard space. We obtained single-trial beta estimates for BOLD response magnitude on each trial by modelling fMRI time series with individual trial regressors (51, 52) using AFNI (53) (onset times for gamble and outcome display and duration modulation with response times or duration on screen respectively). This resulted in 5544 (252/subject) beta estimates at each voxel in the brain, 2772 (126/subject) corresponding to decision and outcome.

### Voxelwise Multilevel Models

We then modeled trial-related estimates of BOLD activity at each voxel in the brain in R (54) (see SI Methods). All p-values are FDR corrected (q=0.05); the cerebellum was excluded from all analyses. Reported coordinates indicate the centre of mass in MNI space and size in number of voxels. Multilevel models had random effects of participant and scanning run included. Linear multilevel models at decision predicted the neural activity with expected gain and loss. Linear multilevel models at outcome predicted neural activity with gain and loss magnitude interacting with choice [coded as 1 (taken), 0 (no response), and −1 (passed)].

To model choices, we computed generalized multilevel models (using a binomial distribution link function) where neural activity at decision predicted choice [coded 1 (taken) and 0 (passed), missed excluded] while controlling for expected gain and loss. Given our interest in trial-related changes, neural activity in choice models was decomposed into three variance components: (1) trial-related variance, beta estimate minus within-scanning run means, (2) scanning run-related variance, within-scanning run means minus within-subjects means, and (3) subject-related variance, within-subjects means minus grand mean.

### ROI Mediation in Multilevel Models

Potential mediating regions were identified by the conjunction of significant effects at decision: (a) expected gain related to increased activity, (b) expected loss related to decreased activity, and (c) trial-related activity predicted responses. Given this conjunction, we performed a liberal cluster correction to exclude small clusters (<15 voxels). For each region that met these criteria, we calculated the within-subjects mean time series across and then used these values to calculate the models necessary for mediation analysis. Mediation analysis in multilevel models presents some unique challenges, for example trial-level mediation effects are nested within person-level effects (40). One could compute separate models for each participant and then average them, but this loses the nested data structure and does not account for individual differences in mediation strength. Thus, we utilized the stacked multilevel regression procedure adapted from recent work exploring mediation analyses in multilevel models (41) to estimate the indirect effect and followed this with Monte Carlo multivariate simulation to estimate confidence limits (41, 42) (SI Methods). This mediation analysis procedure was repeated for each of the identified regions for expected value-choice mediation, and was repeated on pairwise groupings of regions, to explore interregional mediation of choice.

### Mediation Analysis in fMRI

We can be certain about the direction of effects, i.e., brain activity is a product of environmental stimuli and decisions result from brain activity. However, we are less certain about the reliability of these measures. There is substantial measurement error associated with brain activity due to the physics underlying the BOLD response, and this likely results in *underestimation* of the mediation effect (55). In addition, given the number of brain regions or neurons therein, it is likely that variables are omitted. This omission is an unavoidable side-effect of task design, where paradigms must focus on delineating psychologically relevant phenomena. The present approach does not preclude the possibility that activity in other regions might cause both activity in our regions of interest and choices. Omitting a variable may *overestimate* the mediated effect (55). However, this should be explored with separate, hypothesis-driven experiments.

## Acknowledgements

Thank you to Hannah Nohlen, Anthony Romyn, John Tennant, and Thalia Vrantsidis for comments, and to Samantha Mowrer & Amanda Kesek who assisted in earlier stages of the project.

